# Ongoing activation of visual cortex and superior colliculus in the *rd10* mouse model of retinitis pigmentosa

**DOI:** 10.1101/2024.09.02.610761

**Authors:** Thomas Rüland, Kerstin Doerenkamp, Peter Severin Graff, Sophie Wetz, Anoushka Jain, Gerion Nabbefeld, Jana Gehlen, Sara RJ Gilissen, Lutgarde Arckens, Simon Musall, Frank Müller, Björn M. Kampa

**Affiliations:** Molecular and Cellular Physiology (IBI-1), Forschungszentrum Jülich, Germany; Systems Neurophysiology, RWTH Aachen University, Germany; Research Training Group 2416 MultiSenses-MultiScales, RWTH Aachen University, Germany; JARA Brain Institute of Neuroscience and Medicine (INM-10), Forschungszentrum Jülich, Germany; Bioelectronics (IBI-3), Forschungszentrum Jülich, Germany; Research Training Group 2610 InnoRetVision, RWTH Aachen University, Germany; Laboratory of Neuroplasticity and Neuroproteomics -Department of Biology, KU Leuven, Belgium; Leuven Brain Institute, KU Leuven, Belgium

**Keywords:** cross-modal plasticity, oscillatory aberrant activity, retinal degeneration, vision restoration, blindness

## Abstract

Efforts in vision restoration have been focused on a condition called Retinitis Pigmentosa, where photoreceptors in the retina degenerate while the rest of the visual pathway remain mostly intact. Retinal implants that replace the phototransduction process by stimulating retinal ganglion cells have shown promising but limited results in patients so far. Apart from technical limitations, cross-modal plasticity of visual areas might contribute to this problem. We therefore investigated if the primary visual cortex (V1) of the *rd10* mouse model for retinal degeneration became more sensitive to auditory or tactile sensory inputs, potentially hindering retinal stimulation. After reaching complete blindness confirmed by the lack of optomotor responses, activity in visual cortex and superior colliculus (SC) was recorded using Neuropixels probes. While we could not find any significant differences in tactile or auditory responses compared to wildtype mice, the local field potential revealed distinct oscillatory events (0.5 – 6 Hz) in V1 and SC resembling previously observed aberrant activity in the retina of rd10 mice. Further absence of cross-modal plasticity was confirmed by a lacking increase in *zif268* expression in V1 after tactile stimulation. We therefore propose that aberrant retinal activity is transmitted to higher visual areas where it prevents cross-modal changes.

## Introduction

Treatment of blindness is an ongoing and, so far, unsolved endeavor in vision research. As causes are numerous, treatment has focused on a group of diseases called Retinitis Pigmentosa (RP) where patients gradually lose vision as their photoreceptors degrade over time until reaching full blindness (Hartong et al., 2006). While downstream retinal layers do reorganize after longer timescales they generally stay intact in early phases of the disease and therefore provide the unique opportunity to restore vision by selectively replacing the phototransduction process (Jones et al., 2016; Phillips et al., 2010). Different attempts at this have been made using either mostly electrical retinal stimulation, however results have often been limited to the perception of phosphenes, large changes in luminance or high contrast stimuli (Ayton et al., 2014; Nowik et al., 2020; Stingl et al., 2015; Yue et al., 2020).

While technical limitations might be one cause for the apparent lack of success, a different, so far less considered, reason could be cross-modal plasticity leading to reduced activation of former visual areas. As blindness sets in, the visual parts of the brain have been shown to change in function and start processing non-visual information more prominently (Collignon et al., 2008; Merabet and Pascual-Leone, 2010). Different studies found increased activation of visual cortex in blind patients, compared to a normal vision control group, when performing tactile braille reading (Burton et al., 2005; Merabet et al., 2008; Sadato et al., 1996) or auditory localization tasks (Arno et al., 2001; Voss et al., 2008). This phenomenon is thought to be facilitated by different processes that mostly depend on the age at which blindness starts and the investigated visual pathways (Kupers and Ptito, 2014; Mezzera and López-Bendito, 2016). While early loss of vision has been connected to more profound changes in the development of visual pathways (Collignon et al., 2013; Leporé et al., 2010; Noppeney et al., 2005; Pan et al., 2007; Park et al., 2007; Ptito et al., 2008; Shimony et al., 2006), investigations in late blind individuals as well as animal models have shown that changes can happen well after establishment of cortical pathways in adult subjects (Lazzouni et al., 2012; Merabet et al., 2008; Nys et al., 2014; Van Brussel et al., 2011; Voss et al., 2008, 2006). A specific study in enucleated mice showed that even in adult mice, loss of vision can lead to cross-modal changes whereby tactile stimulation also activates parts of the visual cortex (Van Brussel et al., 2011). Depending on the severity and permanence of these changes, they could partially explain the lack of success in vision restoration approaches as visual areas might have lost their sensitivity to the restored visual input.

To address this question, we used the *rd10* mouse model for retinal degeneration which closely mimics the progression of RP and is characterized by a loss of photoreceptors in early adulthood after full formation of the visual system (Chang et al., 2007, 2002, 2000). The progress of the disease in this mouse model was so far mostly characterized by histological investigations of the retina showing slow but progressive photoreceptor degeneration between 3 and 9 weeks of age (Barhoum et al., 2008; Chang et al., 2007; Gargini et al., 2007; Phillips et al., 2010). However, these results did not fully agree with previous behavioral assessments, suggesting that *rd10* mice might lose their visual ability at a later stage in their adulthood (Thomas et al., 2010).

Furthermore, recordings in ex vivo retina preparations revealed aberrant retinal activity which has been discovered most prominently in retinal degeneration models of mouse (Borowska et al., 2011; Margolis et al., 2008; Menzler and Zeck, 2011; Ye and Goo, 2007) but also rat (Sekirnjak et al., 2009) and monkey (Ahn et al., 2022). In the *rd10* mouse model it is characterized by rhythmic events that occur at a frequency of about 3-6 Hz (Biswas et al., 2014; Goo et al., 2011). Evidence suggests that this activity originates in the retina and might be transmitted along the visual pathway to higher visual areas (Dräger and Hubel, 1978; Ivanova et al., 2016). It is however unclear how this activity affects higher visual areas and if it potentially interferes with the processing of incoming sensory input. As the time of vision loss plays an important role in the establishment and severity of cross-modal changes (Voss, 2013) we first investigated the visual ability of *rd10* mice by testing their optomotor response to establish the time point of blindness onset. Subsequently, after allowing sufficient time for potential cross-modal changes to take place, we measured cortical (primary visual cortex, V1) and subcortical (superior colliculus, SC) responses during visual, auditory and tactile stimulation using electrophysiological recordings as well as immediate early gene expression (*zif268*). This aimed to confirm the loss of visual responsiveness in *rd10* mice and test if responses to non-visual stimuli increased in comparison to a normal vision wild type (WT) control group.

As we found no potential change in responses to cross-modal stimuli we assume that the visual cortex remains intact after retinal degeneration. We observed spontaneous oscillations similar to the aberrant retinal activity which might function as continuous activation of downstream visual areas even after loss of visual input. Together, these results provide further insights into the progression and potential treatments of retinal degeneration diseases.

## Materials and Methods

### Optomotor Response (OMR)

#### Experimental Setup

##### Animals

Optomotor response testing was employed in five WT (C57BL/6J, 4 male, 1 female) and six *rd10* (B6.CXB1-Pde6b^rd10^/J, 4 male, 2 female) mice starting at postnatal week 4. Mice were housed in an inverted 12h dark/light cycle and had access to food and water ad libitum. Housing contained no environmental enrichment because of its potential effect on retinal degeneration progression (Barone et al. 2014).

##### Optomotor box and testing

To assess visual ability, mice were placed in a custom-made arena which presented a visual stimulus on four monitors (BenQ GW2270, BenQ, Taiwan) surrounding a raised platform. Square wave gratings with a spatial frequency of 0.05 cycles/deg were presented drifting across the screens with a speed of 33.75 deg/sec. While pre-experiments showed this speed to be best at evoking clear following movements (data not shown) the chosen speed was higher than than the 12 deg/sec used commonly in the literature (Kretschmer et al., 2015, 2013; Prusky et al., 2004). We therefore confirmed that a slower speed (11.25 deg/sec) did not change the acquired results (Fig S1). To achieve different stimulus brightness, the monitor’s background illumination was adjusted and the resulting brightness was measured as the average brightness of black and white bars. Contrast was adjusted by changing the brightness values of black and white bars separately and measuring resulting contrast ratios between the two. Mice were placed on a platform inside of the arena for each stimulus combination (illuminations of 0.2, 1.5, 32, 185 cd/m² at 0.2 and 1.0 contrast levels) for two minutes while their movements were recorded by a camera (Logitech C920, removed IR filter, Logitech, USA) mounted above. For low light conditions, the box was additionally illuminated by an array of infrared LEDs. To ensure that mice were adapted to each light condition tested, a 40 min dark adaptation phase preceded all experiments and mice were subsequently tested in increasingly bright light conditions. Testing was performed consistently at the beginning of the wake cycle (dark-phase) at least 3 times a week.

#### Analysis

Head movement of mice in the optomotor arena was analyzed using the position of both ears as detected by DeepLabCut (Mathis et al., 2018). A line perpendicular to the line connecting both ears was drawn to assess the current head direction of the mouse in each frame. Movement speed of head rotation was then calculated in deg/s over the course of the whole experiment. Visual ability was assessed using a modified OMR ratio (Kretschmer et al., 2015) calculated as follows:

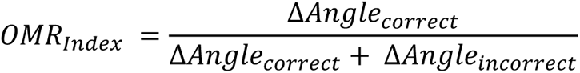

Head movements were labeled as correct or incorrect depending on if they were consistent with the gratings drifting direction. To avoid contamination of the results by behavior unrelated to the stimulus shown, relatively slow (<4 deg/s) and fast movements (>40 deg/s) were excluded from analysis (<2 deg/s and >12 deg/s for slower grating speed used in Fig S1).

OMR ratios shown in Fig 1C were calculated by averaging all OMR ratios acquired in each week in the highest contrast recorded (1.0). Point of vision loss as shown in Fig 1D was calculated by finding the earliest span of three weeks in which the distribution of OMR ratios acquired in each mouse was not different from a distribution with a mean of 0.5 assessed using a Wilcoxon signed rank test (p_adjusted_=0.05, Bonferroni corrected for the number of tests) as implemented in SciPy (Virtanen et al., 2020). Data was handled using pandas dataframes (McKinney, 2010; Reback et al., 2021) and visualized using matplotlib (Hunter, 2007) and seaborn (Waskom, 2021).

**Figure 1.**
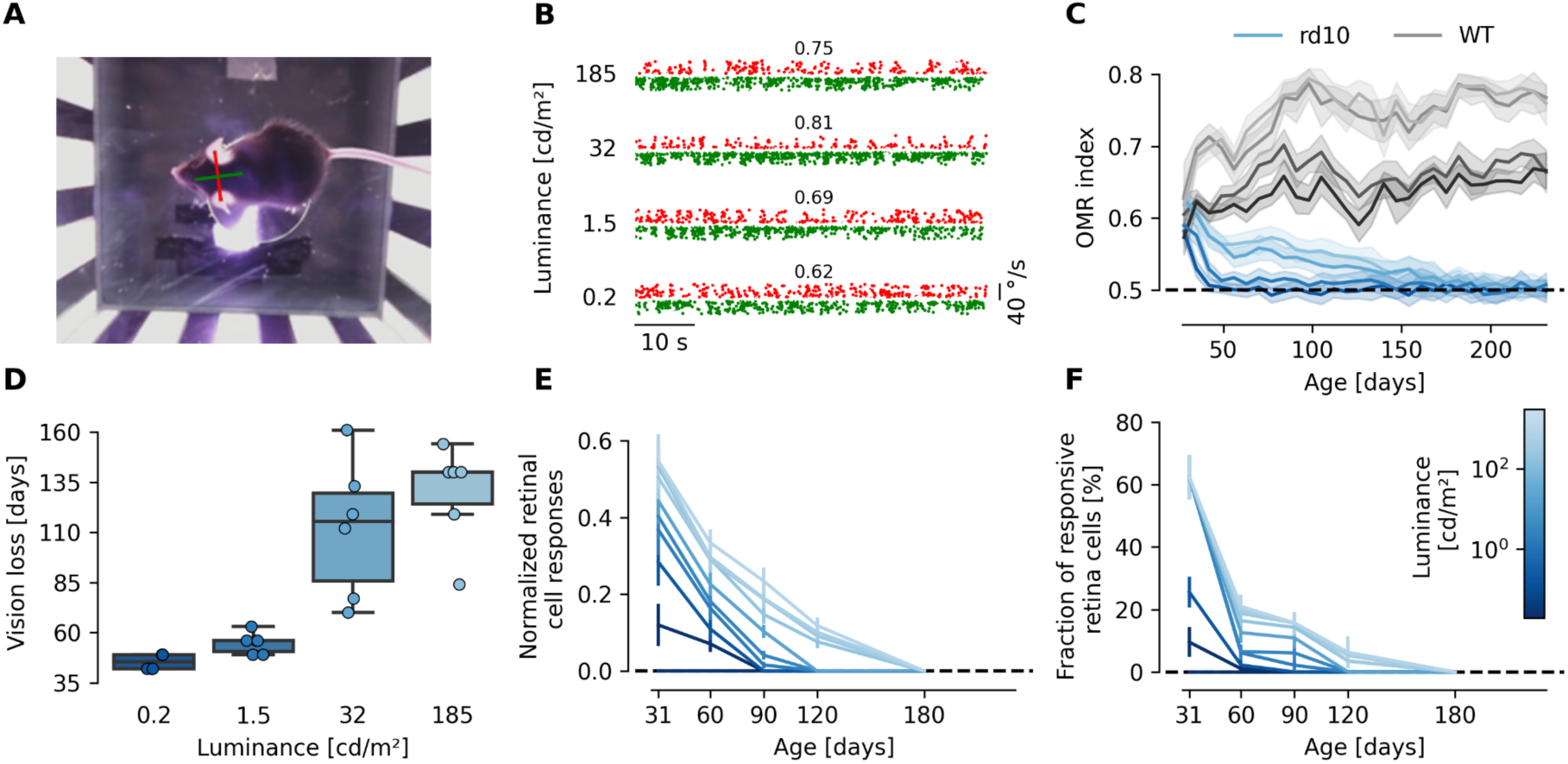
Assessing blindness in *rd10* mice. **(A)** Overhead picture of a mouse in the OMR box which was used to assess head direction. The green line indicates head direction as computed by the perpendicular red line drawn between both ears. **(B)** Examples of head movement speeds acquired at four different illumination strengths. Valid movements along the grating’s movement direction are marked in green, movements opposite to that in red. Ratio of green to red movements increases with increase of the illumination strength (ratios are shown above each trace). **(C)** OMR index indicating stimulus following performance over time for *rd10* (shades of blue) and WT (shades of gray) mice. Illumination strengths are depicted by line brightness, the shaded area depicts the 95% confidence interval. Following performance stays stable for WT mice while *rd10* mice show deteriorating following performance over time (*rd10*: n_mice_=6, n_sessions_=755, WT: n_mice_=5, n_sessions_=523). **(D)** Point of vision loss as determined by a lack of significant following behavior (Wilcoxon signed rank test, p<0.05, Bonferroni adjusted) in each illumination condition for all *rd10* mice tested. Analogue to the time courses shown in **C**, loss of following behavior occurs earliest in lowest illumination and latest in highest. (n_mice_=6) **(E, F)** Comparison of normalized light response and fraction of responsive cells for different age stages as well as illumination strengths in cd/m² (brightness of line). **(E)** Light responses of all remaining ON-cells in extracted retinae of *rd10* mice, normalized to WT responses (P90) for different age stages (WT P90: n_mice_=2, n_cells_=52, *rd10* P31: n_mice_=2, n_cells_=29, P60: n_mice_=6, n_cells_=47, P90: n_mice_=4, n_cells_=41, P120: n_mice_=4, n_cells_=96, P180: n_mice_=2, n_cells_=0). **(F)** Fraction of responsive ON-and OFF-cells in extracted retina of *rd10* mice for different age stages (WT P90: n_mice_=2, n_cells_=178, *rd10* P31: n_mice_=2, n_cells_=214, P60: n_mice_=6, n_cells_=690, P90: n_mice_=5, n_cells_=392, P120: n_mice_=4, n_cells_=373, P180: n_mice_=2, n_cells_=0).

### In Vitro Electrophysiology

#### Experimental Setup

##### Animals

WT animals of the strain C57BL/6 were obtained from Charles Rivers. *Rd10* mice were bred locally from breeding pairs obtained from Jackson (B6.CXB1-*Pde6b^rd10^*/J). In this line, the *rd10* mutation was backcrossed onto the C57BL/6J background for 5 generations before intercrossing to homozygosity. Animals were kept on a 12h light/dark cycle with food and water ad libitum.

##### Multi-electrode-arrays (MEA) recording and electrical stimulation

MEAs containing 60 titanium nitride electrodes (diameter 30 µm, spacing 200 µm, impedance 50 kΩ at 1 kHz) on a glass substrate (Multi Channel Systems MCS GmbH, Reutlingen, Germany) were used. The data acquisition system (MC_Card, Multichannel system, Reutlingen, Germany) consisted of a USB MEA60-Up System, an integrated preamplifier and filter, stimulus generator STG 4002-1.6mA, and a PC. Signals were sampled at 25 kHz/channel.

##### Tissue preparation

Animals were dark-adapted overnight and tissue preparation was performed under dim red light. Briefly, mice were deeply anesthetized with isoflurane and killed by decapitation. Eyeballs were enucleated and retinae isolated. Retinae were cut into two halves and stored in carbonate-buffered AMES solution (pH of ∼7.4), bubbled with 95% O_2_ + 5% CO_2_. For each experiment, one half of the retina was first mounted on a nitrocellulose filter with a hole and then, together with the filter onto the MEA with RGCs towards the electrodes. MEAs were pre-treated in a plasma cleaner (Diener Electronic GmbH + Co. KG, Germany) and coated with 0.5 mg/ml of Poly-D-lysine hydrobromide (PDL, Sigma) overnight. In the recording chamber, the retina was continuously superfused with the carbogenated AMES solution at a flow rate of 3 ml/min at room temperature.

##### Light stimulation

Light stimuli (100 ms long) emitted by a white LED (fig. 2.3.2, ANSI white, 3465K, 185 lm @ 700 mA) below the MEA were triggered by an external stimulator controlled by the software MC-Stimulus. Light intensity started in the scotopic range (equivalent to 0.1 rhodopsin isomerizations per rod outer segment and flash), followed by mesopic and finally photopic stimuli (brightest stimulus 4.18 million rhodopsin isomerizations per rod outer segment and flash).

**Figure 2.**
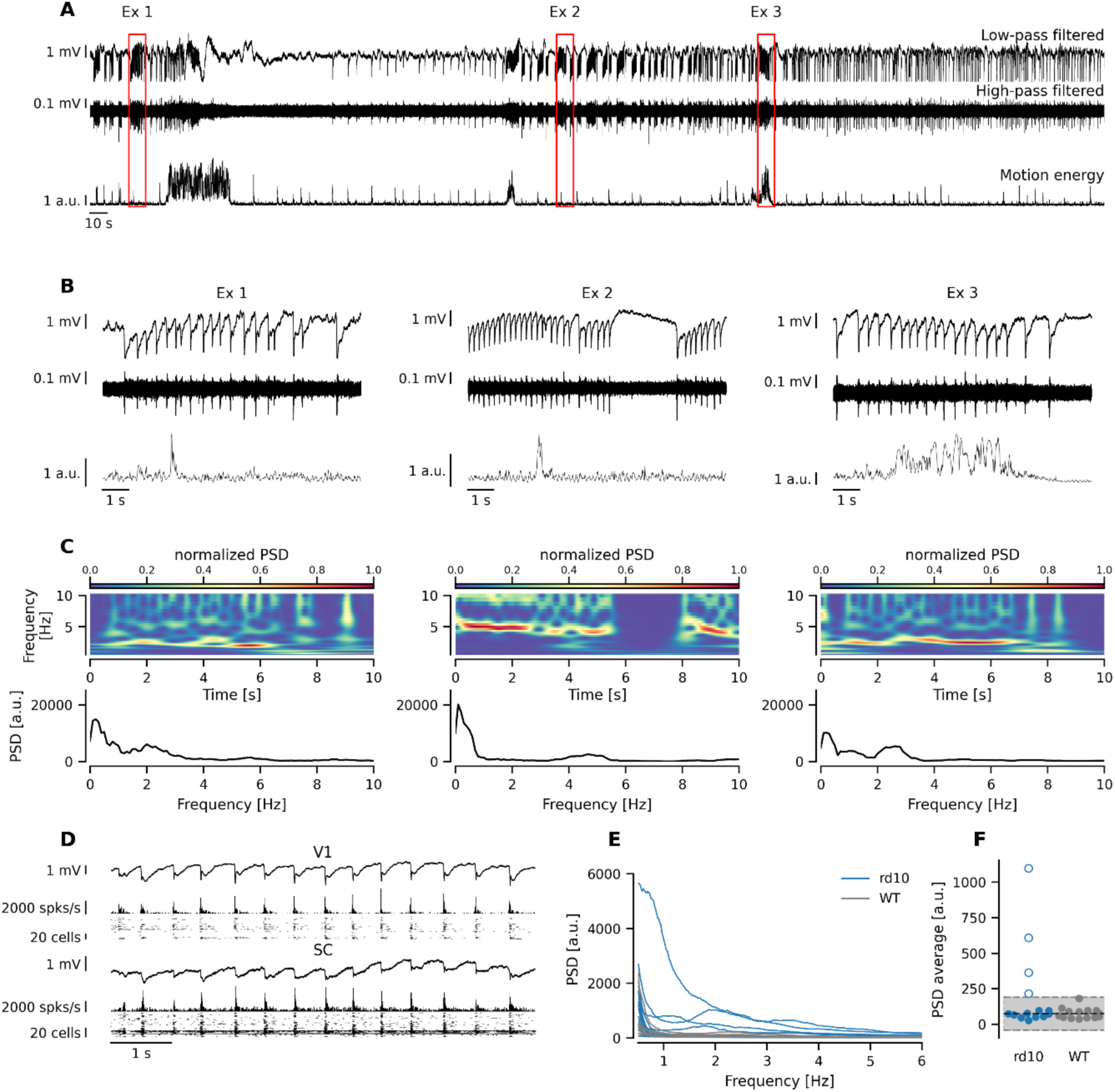
Characterization of aberrant *rd10* activity. **(A)** Excerpt of several minutes of low-pass (**top**) and high-pass (**middle**) filtered *rd10* activity taken from V1. Additionally, the average motion energy of the behavioral cameras is depicted (**bottom**). Red boxes mark 10 s excerpts that are further investigated in (**B**). **(B)** Zoomed in excerpts marked in (**A**) showing oscillatory high amplitude events of varying frequency. Frequency of the events was either relatively stable (Ex 1) or varied over the course of the 10 s excerpts (Ex 2, 3) by changing its timing (Ex 3) or vanishing for short time periods (Ex 2). **(C) Top:** Spectrograms showing the normalized power of frequencies between 0.5 and 10 Hz across the excerpts. **Bottom:** Periodograms showing the power spectral density between 0 and 10 Hz for each of the excerpts shown before. **(D)** Excerpt of an *rd10* recording showing the LFP, the summed up spiking activity and a spike raster plot for V1 and SC simultaneously. Heightened spiking activity can be observed at event peaks in V1 and SC together. **(E)** Periodograms showing the PSD for each recording (WT gray, *rd10* blue) during spontaneous activity. Four *rd10* can be seen to show elevated PSD across the whole spectrum between 0.5 and 6 Hz (rd10: n_mice_=4, n_sessions_=14, WT: n_mice_=4, n_sessions_=14). **(F)** Average PSD between 0.5 and 6 Hz of all recorded sessions for *rd10* and WT mice. Shaded area marks the WT mean (dotted line) ± 3 x standard deviation. Sessions outside of the shaded area (unfilled circles) are found to contain oscillatory activity (*rd10*: n_mice_=4, n_sessions_=14, WT: n_mice_=4, n_sessions_=14).

#### Analysis

Recordings of the individual electrodes were first divided into different units based on the waveform of their action potentials using the software MC_Rack (Multi Channel Systems MCS GmbH, Germany) and offline sorter (Plexon Inc, USA). For quantitative evaluation, light responses were determined by counting action potentials of each ON-GC in a time window of 500 ms after the onset of the light stimulus. For each light intensity, these numbers were averaged for WT and *rd10* retina, respectively, and the averaged values were normalized to the maximal response observed in WT retina.

### In Vivo Electrophysiology

#### Surgery

Surgeries were approved by the State Agency for Nature, Environment, and Consumer Protection (LANUV) of the State of North Rhine-Westphalia (AZ: 84-02.04.2016.A357) and supervised by the local animal welfare office (Institute for Laboratory Animal Science, RWTH Aachen). In preparation for the experiment, mice were implanted with a custom-built headbar and a craniotomy was performed above the visual cortex to allow for Neuropixels probe insertion. For headbar implementation mice were anesthetized using isoflurane inhalation (3% for initiation, 1.5 – 2.5% during the surgery) and placed on a heating pad (TC200 Temperature controller, Thorlabs GmbH, Germany) for body temperature regulation. Analgesia was administered subcutaneously with buprenorphine (0.1 mg/kg Buprenovet, Bayer AG, Germany) for general analgesia as well as bupivacaine (PUREN Pharma GmbH & Co. KG, Germany) for local analgesia at the point of incision. Before incision, hair was removed using hair removal cream (Veet, Reckitt Benckiser Group PLC, United Kingdom) and the skin was disinfected using iodine. An incision was then made at the midline and the skull laying underneath cleaned. Muscles were pushed to the sides and vetbond (3M, USA) was used to fix them in place. A craniotomy of around 1 mm in diameter was then performed above the visual cortex (AP -3.5 mm, ML -2.5mm) and covered with transparent silicone for later recordings. The head bar was placed above the skull and attached using dental cement. For further protection, areas of the skull that were still exposed were covered with vetbond. Post-surgery mice were given buprenorphine (5 ml in 160 ml water) as well as baytril (1 ml/l, Bayer AG; Germany) via their drinking water for 3 days while their recovery was observed. Experimental Setup

##### Animals

Electrophysiological recordings were performed in five WT (C57BL/6J, 4 male, 1 female, between 4 and 11 months old) and four *rd10* (B6.CXB1-Pde6b^rd10^/J, 3 male, 1 female, between 9 and 10 months old) mice at least 10 weeks after blindness onset. Analogue to mice used in the OMR experiments, mice were housed in an inverted 12h dark/light cycle and had access to food and water ad libitum. Housing contained no environmental enrichment because of its potential effect on retinal degeneration progression (Barone et al. 2014).

##### Recordings

To record from V1 and SC simultaneously, mice were head-fixed on a rubber coated wheel and Neuropixels probes (Putzeys et al., 2019) were inserted at an angle between 35° and 38° above V1 (-3.5 ap, -2.5 ml from bregma) 3.6 mm deep into the brain. The bottom 384 channels of the Neuropixels probe were then used to record a high pass filtered action potential signal (0.3 – 10 kHz) and low pass filtered LFP signal (0.5 – 100 Hz) via an external Neuropixels PXIe card (IMEC, Belgium) used in a PXIe-Chassis (PXIe-1071, National Instruments, USA). Triggers for each stimulation type as well as trial start and wheel movements were connected to a BNC breakout board (BNC-2110, National Instruments, USA) and acquired as analogue and digital signals separately using a PXIe card (PXIEe- 8381, National Instruments, USA) in the same PXIe-Chassis and recorded using SpikeGLX (Janelia Farm Research Campus, USA).

##### Stimulation

Visual stimulation was performed using gaussian noise movies displayed at 60 fps (Niell and Stryker, 2008) and was delivered using two 24” Monitors (LG 23MB35PM-B, LG Electronics Deutschland GmbH, Germany) that were placed 18 cm in front of the mouse and rotated by 45° each facing the mouse’s head. Monitor brightness was calibrated to 60 lux. Tactile stimulation consisted of puffs of air that were given in short bursts with a duration of 20 - 50 ms and a pressure of 0.03 bar. Spouts delivering the stimulation were targeted to hit the whiskers of the mouse and aimed so that they did only elicit whisker deflections but no additional startle reactions. For auditory stimulation, one speaker (AST-03208MR-R, Digi-Key Electronics, USA) was placed underneath each monitor and produced clicking sounds at 60 dB tuned for broad auditory stimulation. Experiments were conducted on four consecutive days and started 5-10 minutes after the electrode was inserted to allow for proper settling of the brain tissue around the probe. Stimulation was controlled using a custom MATLAB (3 mice - 3 x *rd10*) or Python (6 mice - 5 x WT, 1 x *rd10*) script and stimuli were delivered ipsi- and contralateral in all possible combinations for 50 trials each. Each trial consisted of three repetitions of the stimulus in question spaced 500 ms apart. The time between individual trials was 3-5 s.

#### Analysis

##### Pre-processing

From the acquired high-pass filtered signal, spikes were extracted and sorted using kilosort 2.5 (Pachitariu et al., 2016). Additional metrics for each unit were computed using the ecephys spike sorting toolbox (Allen Institute for Brain Science) and together with manual unit labels were leveraged to train a custom classifier to exclude noise units. Remaining units were sorted into multi- and single-units by thresholding several quality metrics (amplitude cutoff < 0.1, ISI violations < 0.5, presence ratio > 0.8) and excluding units from analysis that had consistent onset latencies lower than 4 ms to avoid stimulus artifacts. Units were then assigned to different brain areas using their relative depth on the electrode as computed by adapted code from the Cortex-Lab (Steinmetz et al., 2024) and reconstructed probe tracks from the fluorescent traces induced by Dil painted electrodes in the recorded brains. For this, brains were extracted after perfusion and first stored in PFA, then 10% and 30% sucrose solutions sequentially for 24h each. After embedding in Tissue-Tek (Sakura Finetek Europe B.V., Germany), brains were sliced coronally at 100 µm thickness using a cryostat microtome at -20°C and images of each slice acquired using a custom-built fluorescent microscope. Probe tracks were then aligned to the Allen Mouse Common Coordinate Framework (Wang et al., 2020) using the AllenCCF toolbox (Shamash et al., 2018). As it was not always possible to assign each electrode insertion to a specific probe track, histological borders for each area were computed from all four probe tracks and the most conservative estimate used in sensory response analysis. Units were then assigned to each area based on their depth on the probe. For the investigation of the four oscillating *rd10* sessions (Fig 3), the most probable probe was selected for each session and alignment of areas was performed using the electrophysiological alignment GUI by the international brain laboratory (IBL).

**Figure 3.**
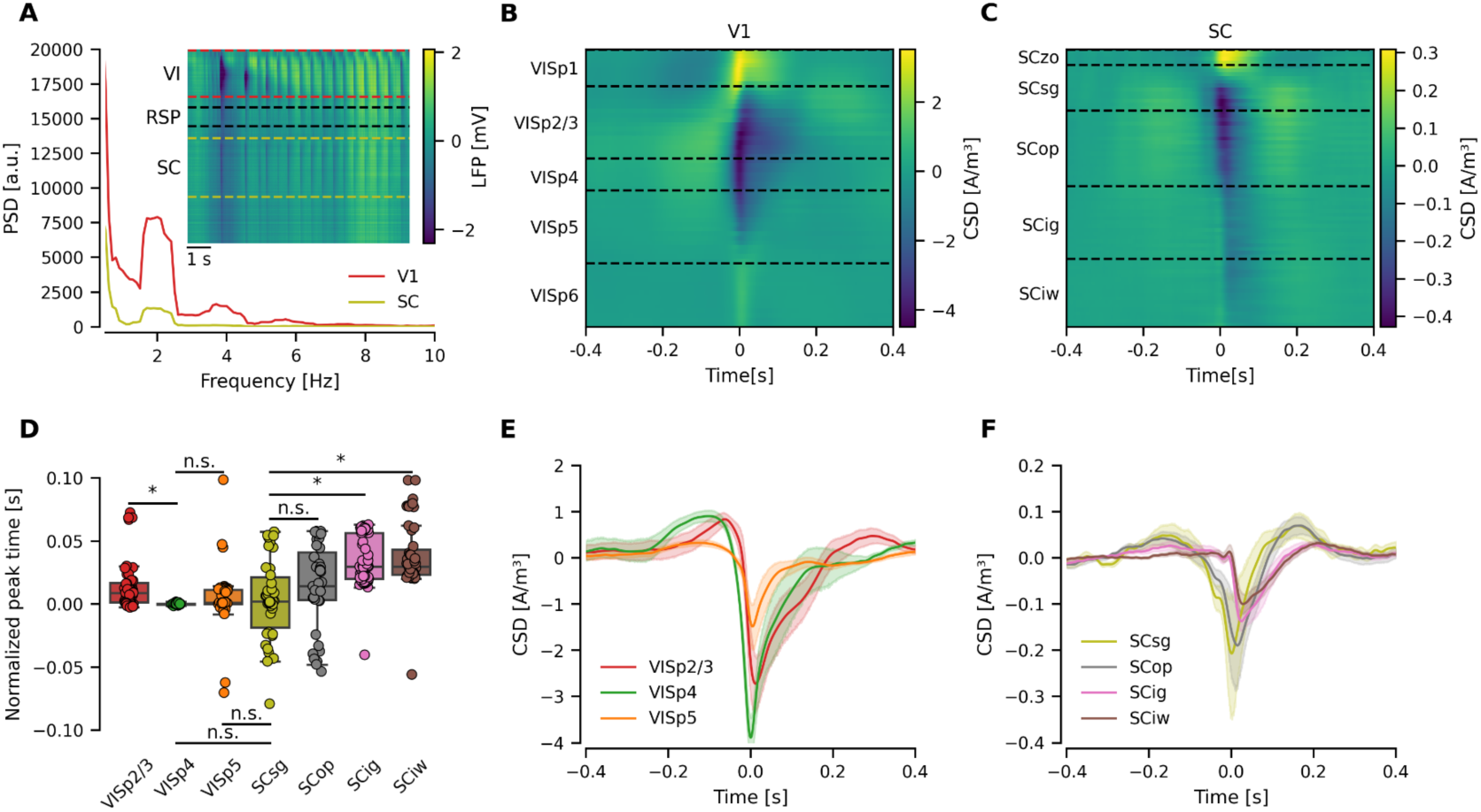
Origin of aberrant activity. **(A)** Periodogram showing the PSD for V1 (red) and SC (yellow) for the excerpt shown in the inset. Inset: Raw LFP of all recorded channels of an *rd10* example recording. Areas identified as V1 (red) and SC (yellow) by histological examination are marked by dashed boxes. V1 and SC show simultaneous oscillations with amplitudes diminishing outside these areas. **(B, C)** Average current source density recorded in oscillating sessions (n_Mice_=4, n_Sessions_=4) 800 ms around a common set of oscillatory event occurrences (manually determined in 500µm depth) for V1 (C) and SC (E). **(D)** Peak times of oscillatory events in all channels for each area recorded (n_Mice_=4, n_Sessions_=4). Pairwise differences were tested for statistical significance using a Dunn-Bonferroni post-hoc-test (p≤0.05) after confirming the existence of statistical differences using a Kruskal-Wallis test (p≤0.05). **(E, F)** CSD averaged for each layer of V1 (D) and SC (F). Shaded area shows the 95% confidence interval (n_Mice_=4, n_Sessions_=4).

##### Sensory responses

For analysis of sensory responses, data from 11 WT sessions (3 mice) and 14 *rd10* sessions (4 mice) was used. For Figure 4, only data excluding the first session of every mouse is displayed to exclude the potential influence of the observed oscillatory activity. Data of all sessions is shown in Supplementary Figure S7. Population responses were computed using all spikes of non-noise units in each session 100 ms before and 100 ms after stimulation onsets. Spikes were binned in 3 ms bins and spike counts were normalized for each session by subtracting the baseline activity (average count 100 ms before stimulus onset) from the resulting signal. Signals were then averaged across sessions and displayed using the 95% percent confidence interval of the bootstrap distribution (Fig 4D, G) as calculated by seaborn 0.11.2 (Waskom, 2021). Comparisons of the average post-stimulus population responses were done by averaging across the normalized spike counts in the 100 ms after stimulus onset and were displayed using boxplots with a maximum whisker length of 1.5 times interquartile range as well as individual data points superimposed. Statistical comparison between WT and *rd10* sessions were done using a Mann-Whitney U test whereas the existence of visual responses was tested using a Wilcoxon signed-rank test. Both were computed in SciPy 1.7.3 (Virtanen et al., 2020) using a significance level of p=0.05.

**Figure 4.**
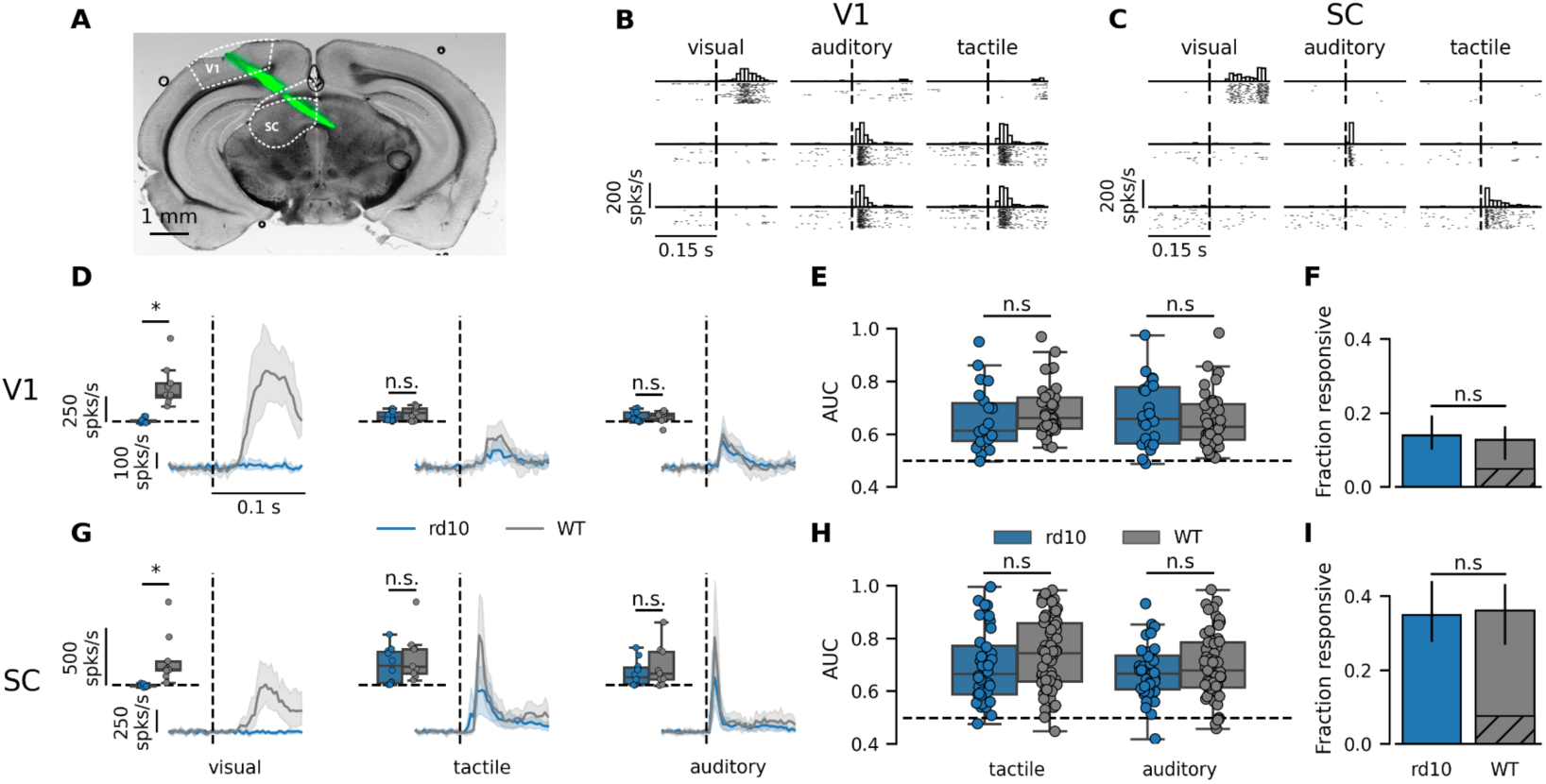
Investigation of cross-modal changes using in vivo electrophysiology. **(A)** Coronal slice of a mouse brain depicting the recording electrode tract (green) as well as the outline of V1 and SC (dotted lines). **(B, C)** PSTH and spike raster plots of three example units in V1 (**B**) and SC (**C**) in response to visual, auditory and tactile stimulation (n_trials_=50). Each row shows the responses of one unit. Stimulus onset is depicted with a dashed line. **(D, G)** Average population responses of all units in *rd10* (blue) and WT (gray) mice in SC (**D**) and V1 (**G**) to each stimulus presented (visual, tactile, auditory). Stimulus onset is marked by the dashed line. Responses were summed up for each session, baseline subtracted and then averaged. Shaded areas indicate the 95% confidence intervals. Boxplots show average population responses for each session (0-100 ms after stimulus onset). Statistical differences were compared using the Mann-Whitney U test (p<0.05) (*rd10*: n_mice_=4, n_sessions_=10, WT: n_mice_=3, n_sessions_=9). **(E, H)** Stimulus separability of single-units that responded to non-visual stimuli were measured using the AUC in *rd10* (blue) and WT (gray) mice depicted as a boxplot and differences measured using Mann-Whitney U test (p≤0.05) (*rd10*: n_mice_=4, V1: n_single-units_=22, SC: n_single-units_=36, WT: n_mice_=3, V1: n_single-units_=39, SC: n_single-units_=61). **(F, I)** Fraction of single-units responsive to auditory or tactile stimulation in SC (**F**) and V1 (Ι) in *rd10* (blue) as well as WT (gray) mice. In WT mice the fraction of single-units that also react to visual stimulation are marked as hatched. Errors are given as 95% confidence intervals as computed by asymptotic normal approximation, differences are compared using Fisher’s exact test (p≤0.05) (*rd10*: n_mice_=4, V1: non-visual: n_single-units_=22, SC: non-visual: n_single-units_=36, WT: n_mice_=3, V1: non-visual: n_single-units_=24, mixed: n_single-units_=15, SC: non-visual: n_single-units_=48, mixed: n_single-units_=13).

Stimulus separability was investigated using the area under the curve (AUC) of a receiver operator characteristic. For each unit, spikes occurring 100 ms before and 100 ms after each stimulation trial were counted and formed two distributions. The detectability of the stimulus based on these two distributions of spikes was measured using the ROC and plotted as hit vs false alarm rate. The area under that curve indicated if both distributions were inseparable (AUC of 0.5) or completely distinct (AUC of 1). Units were marked as significantly responsive when their AUC was higher or as high as at least 99% of AUCs synthetically computed by shuffling pre- and post-trial labels. AUCs of non-visually responsive (responsive to tactile and/or auditory stimulation, non-exclusive) single-units were depicted as boxplot with a maximum whisker length of 1.5 times the interquartile range as well as individual data points superimposed. The amount of non-visually (tactile and/or auditory only) and mixed (responsive to non-visual and visual stimuli) units was determined as described before and divided by the total number of single-units found in the respective region. Fractions were depicted as bar plots and errors are given as 95% confidence intervals as determined by asymptotic normal approximation. A comparison of statistical differences between fractions for each area was assessed using Fisher’s exact test (p=0.05) as implemented in SciPy 1.7.3 (Virtanen et al., 2020). Data was handled using pandas dataframes (McKinney, 2010; Reback et al., 2021) and visualized using matplotlib (Hunter, 2007) and seaborn (Waskom, 2021).

##### Aberrant activity

For analysis of the aberrant activity, the spontaneous activity phases of the low pass filtered signals of 14 WT sessions (4 mice) and 14 *rd10* sessions (4 mice) were used. Only sessions in which at least 10 minutes of continuous spontaneous activity could be measured were included in analysis. As this excluded one of the mice used in the sensory stimulation paradigm from the analysis, we recorded spontaneous activity in two additional WT mice forming the resulting WT group of 4 mice and 14 sessions. Periodograms were computed using the multitaper method as implemented in GhostiPy 0.2.0 (Chu and Kemere, 2021). Smaller excerpts used the full resolution LFP signal (2500 Hz sampling frequency) and a bandwidth of 0.5 Hz (Fig 2C and Fig 3B) whereas periodograms on the complete spontaneous phase of a recording were down sampled to 25 Hz and used a bandwidth of 0.25 Hz (Fig 2E). To determine oscillating sessions, the power spectral density between 0.5 and 6 Hz was integrated and the mean and standard deviation across WT sessions computed. This was used as a baseline and compared to the integrated PSD of *rd10* sessions. When *rd10* sessions exceeded the baseline plus 3 times the standard deviation, they were determined to contain oscillations. Spectrograms that are shown (Fig 2C) were computed using the continuous wavelet transform as implemented by GhostiPy 0.2.0 (Chu and Kemere, 2021) with frequency limits between 0.5 and 10 Hz.

Single oscillatory events were tracked by hand in a recording channel at 500 µm depth from the cortical surface, where amplitudes allowed for clear visibility. The current source density (Fig 3B,C,E,F, Supplementary S3, S4) was calculated using the tracked events as triggers as an inverse current source density as implemented in the py-iCSD (Hagen, 2022) python package and originally developed by Petterson et al. (Pettersen et al., 2006). To compute the average CSD over depth (Fig 3B, C) channels aligned to each layer per recording were upsampled to the maximum number of channels for that layer found in any of the recordings. Each channel was then timeshifted so that the average peak activity in layer VISp4 was centered at event occurrence. The CSD was then averaged for each layer and plotted over depth. For the average comparison per layer (Fig 3E, F) the average CSD per layer was computed for all oscillating recordings. Depictions of the average CSD (Fig 3E, F) show the average per area as a bold line, as well as the 95 % confidence interval as shaded area as computed by seaborn 0.11.2 (Waskom, 2021). Peak times as compared Figure 3D were computed for each channel separately around the occurrence of an oscillatory event, and extracting the peak with highest prominence as determined by SciPy 1.7.3 (Virtanen et al., 2020). Subsequently, channels that had positive peaks and peak times that were not in a 100 ms window around oscillation event occurrence were discarded. A Kruskal-Wallis test was used to check for significant differences between peak-times (p=0.05) and subsequent multiple comparisons between layers were computed using a Dunn-Bonferroni post-hoc-test (p_adjusted_=0.05). Data was handled using pandas dataframes (McKinney, 2010; Reback et al., 2021) and visualized using matplotlib (Hunter, 2007) and seaborn (Waskom, 2021).

##### Motion energy

Behavior videos of two cameras that recorded the mouse from different angles were loaded and encoded as grayscale images using Python. Subsequently, the absolute difference of each pixel across following frames was computed using NumPy (Harris et al., 2020). The mean difference across all pixels between two frames was saved for each timepoint and averaged between both videos recorded to create an “Average motion energy” trace as seen and used in Fig 2B. As one camera failed to record one of the *rd10* oscillating sessions, only the motion energy of one camera was used in Figure S2. As the only moving parts in both videos were corresponding to mouse body, whisker and eye movements, as well as the wheel the mice were running on, the resulting motion energy can be used to approximate the mouse’s behavioral state.

### Immediate Early Gene Expression

#### Experimental Setup

##### Animals

RD10 mice (7 female, 5 male, 9 to 10 months old) and WT controls (6 female, 6 male, 12 to 14 months old) were submitted to three different experimental conditions. After being placed in the dark overnight, they were exposed to either environmental enrichment (via the addition of different toys - EE), environmental enrichment with whisker clipping (WC), or nothing, while remaining in darkness. After a stimulation period of 45 minutes, mice were euthanized with an overdose sodium pentobarbital (Dolethal, 0,2 ml of 60mg/ml i.p.) and brains were carefully removed and snap frozen in 2-methylbutane (Merck, Overijse, Belgium) at -40°C. Brains were stored at -80°C until sectioning. Using a Microm HM 500 OM cryostat (Walldorf, Germany), 25 µm-thick coronal sections were cut and collected on SuperFrost Plus Adhesion slides (10149870, Thermo Fisher Scientific). Sections were stored at -20 °C until the start of the HCR.

##### Hybridization chain reaction

Probe pairs were generated with in situ_probe_generator (Null and Özpalat, 2020), validated for specificity using Blastn (www.NCBI.com), and ordered from Integrated DNA Technologies, Inc (IDT). Third generation in situ hybridization chain reaction (HCR v3.0 ; Choi et al., 2018) was adapted for cryosections (Van houcke et al., 2021). Cryosections were fixated for 30’ minutes in 4% Paraformaldehyde and dehydrated via an increasing alcohol series (50% - 70% - 100% - 100%). Next, sections were washed in TBS (0,3% Triton-X-100 in PBS), two times in PBS-DEPC and once in 5x SSCT (0.1% Tween-20 in saline-sodium citrate buffer (SSC)), each time for 10 minutes. All steps of the hybridization process were performed in a humidity chamber. Probe-hybridization buffer (30% formamide, 25% 5x SSC, 9 mM citric acid pH 6, 0,1% Tween 20, 50 ug/ml heparin, 1 x Denhardts solution, 10% dextran solution; fill up to 40 ml with ultrapure H_2_O) was added to each glass slide (800µl) and placed at 37°C for 30 minutes. After removal of the probe-hybridization buffer, 75µl of probe solution (0.3 pmol of each probe in 75µl probe-hybridization buffer) was added on top of the glass slide and incubated overnight (12-16h) at 37°C. The next day, excess probe was removed by subsequent rinsing with 25% 5X SSCT, 50% 5X SSCT, 75% SSCT and twice 100% 5X SSCT, each time for 15 minutes. To amplify the signal, 800µl of amplification buffer (5 x SSC, 0,1% Tween 20, 10% Dextran sulfate, fill up to 40 ml with ultrapure H_2_O) was added to the glass slides for at least 30 min at RT. After removal of the amplification buffer, sections were incubated overnight (12-16h) with 75µl of the hairpin solution (Snap-cooled hairpins conjugated to a 546 fluorophore in amplification buffer, 9pmol per glass slide or 4,5pmol for each hairpin) per glass slide in the dark at room temperature. After hybridization and amplification steps, the slides were washed three times in 5x SSCT for 10 minutes, followed by Dapi counterstaining (SSCT-DAPI, 1/1000) for 20 minutes and mounting.

#### Analysis

All HCR related images have been generated with a Zeiss light microscope (Zeiss Axio Imager Z1) equipped with an AxioCam MRm camera (1388 x 1040 px) at 20x. Images were analyzed with Image J by measuring the mean gray value. For pictures of the visual cortex, all cortical layers were measured, while for pictures of the somatosensory cortex, only layer 4 (the barrels) was analyzed since only the whiskers were manipulated and there was no full removal of all somatosensory input. One-Way ANOVA (p=0.05) (Fig S8) and Two-Way ANOVA (p=0.05) (Fig 5) were used for assessment of statistical difference whereas single groups were subsequently compared using the Tukey’s multiple comparisons test (p_adjusted_=0.05).

**Figure 5.**
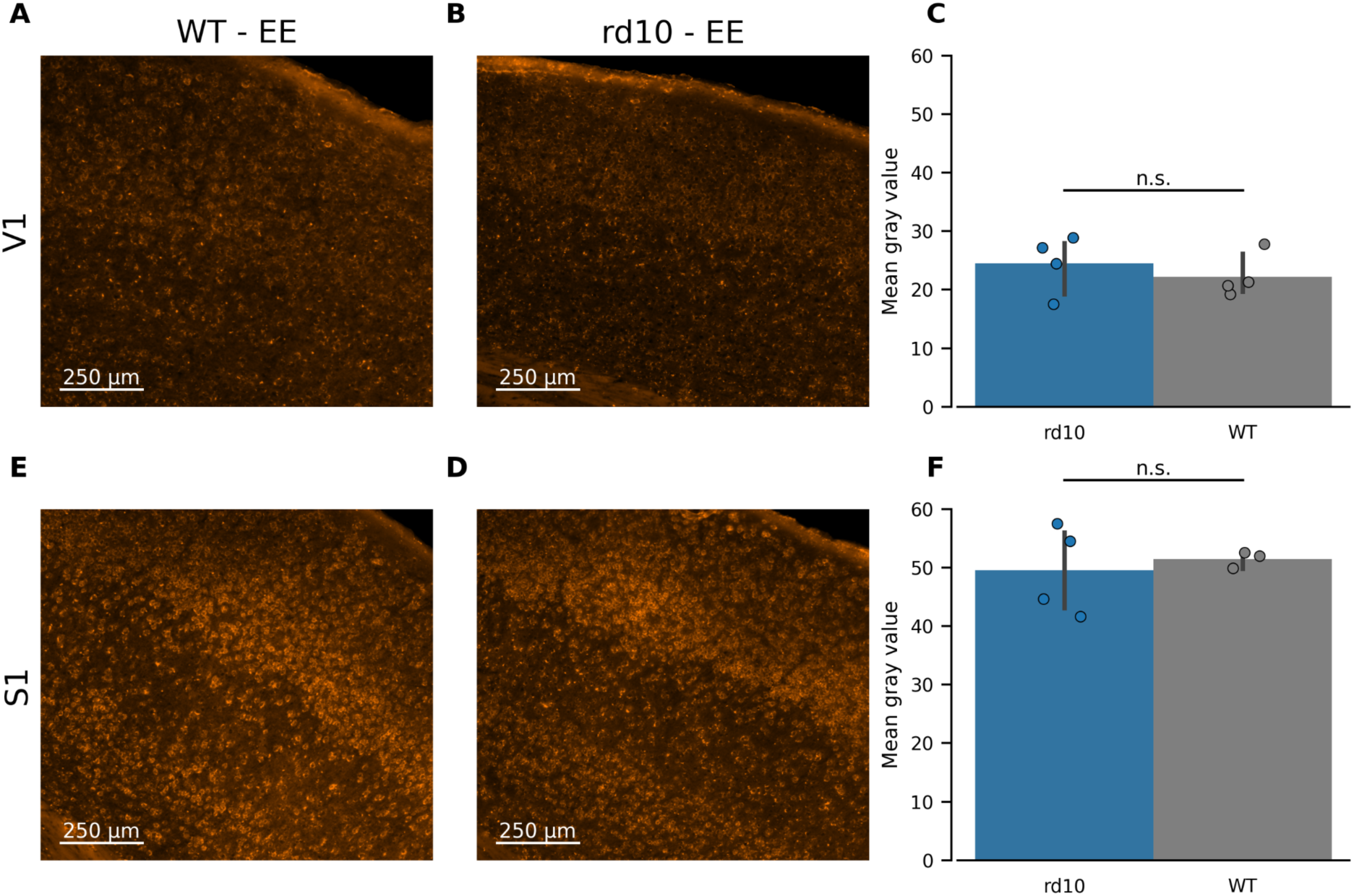
Investigation of cross-modal changes using immediate early gene expression (*zif268*). Example field of views showing the visual cortex **(A, B)** and somatosensory cortex **(E, D)** for WT **(A, E)** and *rd10* **(B, D)** mice exposed to an enriched environment. **(C, F)** Comparison of the average *zif268* expression levels in V1 **(C)** and the barrel field in somatosensory cortex **(F)** after tactile stimulation between WT mice and *rd10* mice. Differences were tested for differences using a Two-Way Anova (p≤0.05) and a subsequent Tukey’s multiple comparison test (p_adjusted_≤0.05) (*rd10 V1*: n_mice_=4, *S1*: n_mice_=4, WT V1: n_mice_=4, S1: n_mice_=3).

## Results

### Progressive loss of vision in rd10 mice

To study potential changes in the visual brain areas of *rd10* mice, we first assessed their degrading visual ability using the optomotor response (OMR). This test allows determining the onset of blindness and to compare our results with previously described changes in relation to this onset (Van Brussel et al., 2011; Voss, 2013). Using four different illumination conditions we measured the OMR of *rd10* as well as a regularly seeing WT control group in response to drifting square wave gratings during the progression of the disease. Mice were situated in an arena where they were presented with a moving grating pattern on surrounding monitors and their head direction was tracked with an overhead camera (Fig 1A). To determine remaining vision, we used the modified OMR index (Kretschmer et al., 2015) which quantifies head movements of the mouse, following and opposite to the grating drifting direction (Fig 1B). It was apparent that higher illumination conditions also produced higher OMR indices as a larger fraction of head movements was related to the presented stimulus. The development of the ratio showed that WT mice improved their stimulus following behavior during the first days of the experiments until a stable performance was reached for all illuminations. Conversely, *rd10* mice, while starting at comparable OMR indices as WT mice, showed a stark decline in stimulus following behavior (Fig 1C). This decrease was observed earlier and more pronounced in low illumination conditions (0.2, 1.5 cd/m²) but also manifested in higher illumination conditions (32, 185 cd/m²), albeit progressing slower. Ultimately, *rd10* mice reached a baseline OMR index of 0.5 in all illumination conditions, demonstrating that no meaningful following behavior was observable anymore. We quantified this time point for each mouse separately and showed that in low light conditions, vision was lost on average in week 6.5 ± 4.9 whereas in the highest light condition mice continued to show following behavior up until week 18.5 ± 3.6 (Fig 1D). Given that previous histological investigations had shown maximum photoreceptor loss at postnatal day P25 (Barhoum et al., 2008) and complete loss until P60 (Gargini et al., 2007) this was rather surprising. We therefore recorded the electrical response of ganglion cells in extracted retinae of *rd10* to bright light flashes to confirm our behavioral results (Fig 1E, F). Here, we could show that at high luminance levels of the visual stimulation (≈190 cd/m²), vastly reduced responses remained visible up until P120 in a small population of cells (Fig 1E, F). Only at P180 we were not able to detect any remaining activity in response to the shown flashes. We therefore performed investigations of potential cortical changes at least 10 weeks later to ensure that cortical changes had ample time to develop, as suggested by investigations of enucleated mice beforehand (Van Brussel et al., 2011).

### Aberrant oscillatory activity in visual areas of *rd10* mice

After complete blindness onset, we waited for about 10 weeks to give *rd10* mice ample time for changes in the visual system to take hold. Subsequently, we started the examination of neuronal activity using high-density Neuropixels probes to record the electrical activity of the visual cortex and SC simultaneously. When electrophysiological recordings were performed, prominent oscillatory events were observed in all *rd10* mice on the first day of recording. This activity manifested in the low pass local field potential (LFP) as well as high pass filtered spiking as a series of high amplitude events with varying frequency typically between 0.5 and 6 Hz (Fig 2A-C). Spontaneous oscillations occurred randomly for brief periods and changed their frequency over the course of smaller bouts (Fig 2A). Animal motion seemed to have a modulatory influence on frequency of observed oscillations (Fig 2B, Ex 3, Fig S1) but was not necessary for the occurrence of oscillatory events. Oscillatory events were most prominent in V1 but could also be observed in SC where event amplitudes were typically diminished (Fig 2D, 3A). However, when events were visible, they co-occurred with simultaneous spiking activity across a large number of neurons in both areas (Fig 2D). To quantify the occurrence of oscillations, the power spectral density (PSD) of spontaneous activity in each session was plotted (Fig 2E) and quantified (Fig 2G). We found a clear increase in a subset of *rd10* sessions which corresponded to the first session of each mouse in which events were also clearly visible by eye (Fig 2F). These sessions exceeded the average WT PSD between 0.5 and 6 Hz by more than three standard deviations (75.6 ± 3 x 38.8, shaded area Fig 2F) and were subsequently identified as oscillating (unfilled circles in the integrated PSD, Fig 2F).

### Aberrant activity might have retinal origin

When aberrant activity was visible in V1 and SC simultaneously, the frequency of both signals matched closely as shown in an example excerpt (Fig 3A). Amplitudes of the signal were diminishing over the depth of each area (Fig 3A inset) and generally seemed to be lower in SC compared to V1 (Fig 3E, F). The aberrant activity furthermore seemed to subside in the void space between retrosplenial cortex (RSP, Fig 3A inset) and SC hinting at a physiological occurrence opposed to an artificial source. Due to the separate but simultaneous occurrence of aberrant activity in both areas we suspected that a common external source might be the driving force behind it. While a visual origin seemed to be the most obvious common input to V1 and SC, projections of other sensory areas are known to terminate in V1 and especially lower SC layers (Dräger and Hubel, 1976; Ito and Feldheim, 2018; Triplett et al., 2012; Zingg et al., 2017). We therefore computed the current source density (CDS) using the LFP recorded in V1 and SC to determine the most probable origin of the oscillatory events.

The average CSD in V1 and SC revealed that both areas showed a strong negative CSD in layers VISp2/3, VISp4 and VISp5 of V1 (Fig 3B) as well as SCsg and SCop of SC (Fig 3C). As a negative CSD is associated with activation of the surrounding area, we assumed that these layers are the most probable candidates for the signal source. To determine if there is a clear sequential order of the layer activation we further investigated the relative timing of the CSD peaks across these layers (Fig 3D-F). In V1, the average CSD showed the earliest peak times in VISp4 (0.2 ± 0.8 ms), followed by layers VISp5 (4.5 ± 23.6 ms, p_vsVISp4_=1) and VISp2/3 (15.0 ± 19.8 ms, p_vsVISp4_=0.013) (Fig 3D). Whereas in SC the earliest peak was visible in SCsg (3.7 ± 31.7 ms) being followed by deeper layers SCop (14.3 ± 33.5 ms, p_vsSCsg_=0.57), SCig (33.7 ± 19.1 ms, p_vsSCsg_=7.85 x 10^-8^) and SCiw (38.0 ± 26.6 ms, p_vsSCsg_=3.5 x 10^-8^, Kruskal-Wallis and Dunn-Bonferroni post-hoc-test) (Fig 3F). Both areas therefore showed the earliest activation in layers which are predominantly receiving visual inputs, with layer VISp4 and VISp5 receiving input indirectly from retina via the lateral geniculate nucleus (LGN) and layer SCsg/SCop directly from retina as well as through feedback from V1 (Ito and Feldheim, 2018). We could observe the same relationship in all four sessions recorded in both V1 and SC (Supplementary Figures S5, S6). Furthermore, there was no significant difference between the peak times of VISp5 and SCsg (p=1) suggesting that oscillations in SC were not solely driven by V1 to SC transmission. We therefore hypothesized that the aberrant oscillatory activity enters V1 and SC separately through their visual input layers and then spreads across each area. While this does not fully confirm a retinal origin of the signal, these results strongly suggest a source higher up the visual processing stream.

### Lack of cross-modal sensory enhancements in *rd10* visual cortex and SC

While the existence of aberrant oscillatory activity in the visual cortex and SC of *rd10* mice could be established, its effect on neuronal processing in both areas remains unknown. Due to ongoing attempts to stimulate the degenerated retina for vision restoration therapies, the ability of central areas to process visually evoked stimuli after blindness onset is of great interest. However, a potential caveat could be that visual areas can become sensitive to other modalities like sound or touch. We therefore investigated whether V1 and SC showed stable responses to non-visual (auditory and tactile) stimulation and whether these responses were significantly altered in comparison to a regularly seeing WT control group. We recorded responses using Neuropixels probes in V1 and SC simultaneously (Fig 4A). To establish the general efficiency of the chosen stimulation, responses of example units in V1 and SC of *rd10* and WT mice, showing reliable responses to each type of stimulation across trials, are shown in Figure 4B and C.

As we could not rule out that the observed aberrant oscillatory activity had an influence on stimulus responses due to masking effects, we compared the non-visual responsiveness of previously determined oscillating to non-oscillating sessions and WT sessions (Supplementary Fig S7). While we could not see differences in population stimulus responses, there was a significant reduction of auditory responsiveness of single units in SC in oscillating sessions (0.57 ± 0.12) compared to non-oscillating sessions (0.67 ± 0.10, p=0.021) and WT sessions (0.69 ± 0.13, p=0.014) as well as tactile responses in V1 in oscillating sessions (0.58 ± 0.04) compared to WT sessions (0.69 ± 0.10, p=0.018). We therefore decided to exclude oscillating sessions from subsequent sensory analysis to avoid potential masking effects. As oscillations regularly occurred on the first day of recording, we also excluded the first day of recording for all WT sessions to exclude a potential influence of the recording day.

For a first comparison of rd10 and WT responses, the baseline subtracted summed-up population response to each stimulation type is shown for V1 and SC in rd10 and WT mice (Fig 4D, G). While WT mice showed strong neural responses to visual stimulation in V1 (367 ± 219 spikes/s, p_vsVis_=0.004) and SC (224 ± 224 spikes/s, p_vsVis_=0.004), these were completely absent in *rd10* mice (V1: 6 ± 25 spikes/s, p_vsVis_=0.77, SC: -4 ± 12 spikes/s, p_vsVis_=0.43, Wilcoxon signed rank test) aligning with our OMR and retinal activation investigations (Fig 1). Analyzing responses to non-visual stimulation (auditory and tactile) in V1 and SC revealed that responses to both were similar and not significantly different in *rd10* and WT mice (V1: rd10_tac_ 53 ± 50 spikes/s, WT_tac_ 79 ± 69 spikes/s, p_tac_=0.54, rd10_aud_ 49 ± 55 spikes /s, WT_aud_ 46 ± 63 spikes/s, p_aud_=0.90, SC: rd10_tac_ 174 ± 162 spikes /s, WT_tac_ 251 ± 216 spikes /s p_tac_=0.31, rd10_aud_ 105 ± 117 spikes /s, WT_aud_ 166 ± 187 spikes/s, p_aud_=0.60 Mann-Whitney U). We then looked at potential differences on a single-unit level. We used stimulus separability by quantifying the area under the curve (AUC) of a receiver operator characteristic of single unit responses to tactile and auditory stimuli. As no single-units were found with significant responses to visual stimulation (data not shown), we focused on single-units that were responsive to auditory or tactile stimulation. Comparing the AUC between *rd10* and WT mice in V1 and SC for both non-visual stimulations (Fig 4E, H) did not show any significant difference between the two mouse groups (V1: rd10_tac_ 0.66 ± 0.12, WT_tac_ 0.69 ± 0.10, p_tac_=0.12, rd10_aud_ 0.68 ± 0.13, WT_aud_ 0.65 ± 0.10, p_aud_=0.55, SC: rd10_tac_ 0.70 ± 0.14, WT_tac_ 0.74 ± 0.14, p_tac_=0.07, rd10_aud_ 0.67 ± 0.10, WT_aud_ 0.70 ± 0.13, p_aud_=0.39 Mann-Whitney U). Furthermore, quantifying the fraction of neurons in V1 and SC that responded to non-visual stimulation in each mouse group (Fig 4F, I) also showed no significant differences (V1: p=0.77, SC: p=0.90, Fisher’s exact test). In summary, we found no evidence for cross-modal plasticity processes in the visual areas of *rd10* mice, even when those were given ample time to develop after blindness onset.

### Lack of increased immediate gene expression in *rd10* visual cortex in response to tactile stimulation

To further confirm that the lack of changes in response to non-visual stimulation in the electrophysiological recordings was not caused by an interference from undetected aberrant oscillatory activity, we investigated the general activity levels of the visual and somatosensory cortex using *zif268* expression in fully blind *rd10* mice after tactile stimulation. Previous studies had shown that adult mice that were visually deprived showed a clear increase in visual cortex responses to tactile stimulation compared to a normally seeing control group (Van Brussel et al., 2011). We therefore used a similar setup in which *rd10* and WT were kept in darkness and received tactile stimulation with intact or cut whiskers. Tactile stimulation consisted of several novel objects that were placed in the cage for an hour for the animals to explore. We found that this procedure did indeed raise activity in the somatosensory cortex of WT and *rd10* mice, confirming its efficiency as tactile stimulation (WT - WC: 35.11 ± 2.80, rd10 - EE: 49.54 ± 7.63 p=0.021, WT - EE: 51.45 ± 1.40, p=0.016) (Figure S8 Supplementary Material). We then compared activity levels in the visual cortex as well as the barrel field of somatosensory cortex after tactile stimulation between *rd10* and WT mice. Here we saw no difference in activity levels between tactile stimulated *rd10* and WT mice neither in visual cortex (rd10 - EE 24.49 ± 4.99, WT - EE 22.23 ± 3.79, p=0.925) (Figure 5C) nor in somatosensory cortex (rd10 - EE 49.54 ± 7.63, WT - EE 51.45 ± 1.40, p=0.993) (Figure 5F). This confirmed the results acquired in the electrophysiological investigation before and further solidifies the lack of cross-modal changes after blindness induced by retinal degeneration in the *rd10* mouse model.

## Discussion

We could show that behavioral progression of vision loss in the *rd10* mouse model is much slower than previously suggested by histological investigations. Our results showed that despite the maximal loss of photoreceptors being reported around postnatal day 25 (P25) (Barhoum et al., 2008), vision declines for several weeks after that and only completely subsides in bright light conditions at around week 20. We confirmed these findings by performing electrophysiological MEA recordings in extracted *rd10* retinae revealing a small fraction of retinal ganglion cells that are active during stimulation with high light intensities at P120 and a lack of responses in P180. While remnants of functional photoreceptors have been found in histological investigations for up to 9 months, their behavioral relevance was highly questionable so far (Barone et al., 2014; Gargini et al., 2007). Together with the OMR findings by Thomas et al. (Thomas et al., 2010) that showed a decrease in visual ability of *rd10* mice over a time span of around 6 months, our results indicate that some form of vision remains functional in the *rd10* mouse model over much longer timescales than previously expected.

This prompted our further investigation of cross-modal changes to be conducted at least 10 weeks after complete blindness onset as previous studies using immediate early gene expression in mice had indicated full reactivation of visual cortex after enucleation by then (Van Brussel et al., 2011). We investigated potential cross-modal changes using the same approach but found no evidence for stronger activation in the visual cortex of *rd10* mice in response to tactile stimulation compared to a WT control group. The same was true when we measured activity on the population as well as the single-unit level in V1 and SC using Neuropixels probes. We found no significant differences in population responses to non-visual stimulation between *rd10* and WT mice. Furthermore, neither single-unit response amplitude nor the fraction of stimulus-responsive single-units were statistically different. Similar evidence was found in our recent study that investigated electrophysiological properties of pyramidal neurons ex vivo in the visual cortex of *rd10* mice (Halfmann et al., 2023). Only minor differences were found during peak retina degeneration but not at later stages, reinforcing the idea of a largely unchanged cortical landscape in the *rd10* mouse model. While this confirmed our acquired results, it was rather surprising given that previous evidence pointed towards cortical changes in response to vision loss in other rodent mouse models (Nys et al., 2014; Piché et al., 2007; Van Brussel et al., 2011; Voss et al., 2006).

The low pass filtered part (LFP) of the electrophysiologically acquired in vivo data revealed that *rd10* mice showed prominent oscillatory events in visual cortex and SC that were occurring rhythmically at frequencies changing between 0.5 and 6 Hz. While their strength and quantity varied greatly between sessions, these oscillatory events showed striking similarity with aberrant activity that has been recorded in degenerated retinas before (Biswas et al., 2014; Gehlen et al., 2020; Goo et al., 2011; Ivanova et al., 2016). This was consistent with research conducted in *rd1* and *rd10* mice in cortex and SC that separately found similar rhythmic activity (Dräger and Hubel, 1978; Ivanova et al., 2016). While the observed activity was modulated by the movements of the mouse (Fig S2, Fig 2A, B), the occurrence of oscillatory events was not dependent on animal movements and long periods of oscillations could be observed also during resting phases (Fig 2A). Events were visible in V1 and to a lower extent in SC separately in the LFP as well as the spiking activity. Calculating the CSD for each area separately showed that oscillation onsets were earliest in layers receiving visual input either from the LGN (VISp4, VISp5) or directly from the retina or indirectly via feedback projections from V1 (SCsg, SCop). This effect was visible in the average as well as each individual session CSD (Fig 3C-F, Supplementary Figure S3-6). While we cannot rule out that the observed oscillatory activity has its origin somewhere along the visual pathway, its simultaneous occurrence in V1 and SC either hints at a common origin at the retinal level or a transmission between the two regions. Furthermore, no significant difference in earliest activity in visual input layer VISp4 and SCsg (p=0.47) as well as visual output layer VISp5 and earliest activity SCsg (p=1) could be found. We therefore could not establish a clear directionality between the two areas. Together with the previous work in the field we however suspect that observed cortical oscillations are either directly transmitted or indirectly caused by aberrant retinal activity of the *rd10* mouse model.

This view is reinforced by a study by Dräger and Hubel that saw aberrant activity in the cortex of retina degenerated mice that could be abolished and reintroduced by asphyxiation of the eye (Dräger and Hubel, 1978), strongly hinting at a retinal origin of the signal. How exactly the aberrant activity is propagated from retina to V1 and SC however, is still unclear. Separate transmission from the retina to both areas appears to be the most parsimonious possibility. Yet, a clear lead of SC (which is directly connected to the retina compared to V1) could not be observed. Furthermore, SC showed lower event amplitudes and spiking occurrence (Fig 2D, 3A, 3F), hinting at a more prominent role of V1 in the processing of the oscillatory signal.

If we assume that the aberrant oscillatory activity observed in downstream visual pathway areas is of retinal origin, this could explain the lack of cross-modal changes in this retinal degeneration mouse model. As spiking activity in both V1 and SC was highly linked to the onset times of oscillating events, one possibility would be that oscillatory events might mask sensory responses to non-visual stimuli leading to underestimation of the cross-modal stimulus responses. To exclude that possibility, we excluded sessions with obvious oscillations from sensory response analysis and did not see any significant changes in the remaining responses. Additionally, we investigated general activity levels in V1 and S1 of *rd10* and WT mice using immediate early gene expression. As this should not be affected by stimulus masking we expected general changes in activity to become visible here. However, results did not show a general rise in activity levels in response to tactile stimulation, further enhancing the apparent lack of changes to the response profile (Fig 5).

We therefore alternatively propose that the observed sustained activation of higher visual areas by aberrant activity might prevent the induction of cross-modal plasticity. Full blindness onset in *rd10* mice occurs after the formation of the visual system as evident by our and previous OMR and ERG recordings (Chang et al., 2007; Thomas et al., 2010). Therefore, potential cross-modal changes are expected to be realized by upscaling of existing connections between visual and non-visual sensory areas due to a lack of primary sensory (visual) input (Merabet and Pascual-Leone, 2010). Assuming that aberrant retinal activity is strong enough to drive spiking activity in visual brain areas, upscaling of connections to non-visual areas in response to lacking input would be prevented.

This lack of cross-modal changes in the *rd10* mouse model together with its clinically relevant phenotype, make it a prime candidate for the further investigation of vision restoration techniques. If functionality of the visual brain areas as well as downstream retinal circuits are conserved, retinal (Ayton et al., 2020; Hadjinicolaou et al., 2012; Hallum and Dakin, 2021; Humayun et al., 2012; Keserü et al., 2012; Klauke et al., 2011; Mathieson et al., 2012; Rizzo, 2011; Stingl et al., 2013; Zrenner et al., 2011) as well as cortical implants (Dobelle, 2000; Normann et al., 2009) could be employed in the mouse model to screen for most promising approaches. While retinal implants in particular have only been sparsely used in animal models so far (Alteheld et al., 2007; Eckhorn et al., 2004; Gekeler et al., 2004; Lohmann et al., 2017), other techniques like retinal sheet transplantation (Foik et al., 2018; Lin et al., 2018), expression of channelrhodopsin in ganglion cells (Reh et al., 2021) as well as replacement of photoreceptors using photovoltaic nanowires arrays (Tang et al., 2018) were successfully applied in rodents and could strengthen the *rd10* mouse models position as a valuable model organism for vision restoration research.

Our findings might also shed more light on possible treatment options of retinitis pigmentosa (RP) in humans. While rhythmic retinal activity is a widespread phenomenon of retinal degeneration and has been described most prominently in the rd mouse model (Borowska et al., 2011; Margolis et al., 2008; Menzler and Zeck, 2011; Ye and Goo, 2007) and to a lesser extent also in retinally degenerated rats (Sekirnjak et al., 2009) and recently monkeys (Ahn et al., 2022), no proof of its occurrence in RP patients has been found so far. Furthermore, cross-modal changes have been found in several studies that also included RP patients, (Burton et al., 2005, 2002; Sadato et al., 2002) hinting at an absence of retinal aberrant activity or at a decrease of its proposed neuroprotective effect. If aberrant activity is indeed absent, artificially stimulating the retina of patients suffering from retinitis pigmentosa during the progression of the disease might induce a similar protective effect as the proposed transmittance of aberrant retinal activity and could help in keeping the visual cortex receptive to visual inputs. Artificial stimulation of the eyes of affected individuals has been performed before using transcorneal electrical stimulation (TES) and saw positive effects on slowing down the disease’s progression (Schatz et al., 2011; Sinim Kahraman and Oner, 2020). While its function was mostly seen in its neuroprotective function of retinal cells as well as the optical nerve, our results suggest that a positive effect might similarly be achieved for downstream visual areas. This would further enhance the technique’s importance as a possible treatment option for patients that suffer from retinitis pigmentosa and indicate that an early treatment start might offer additional advantages.

## Acknowledgements

The authors would like to thank Dr. Christopher Wiesbrock for his help in the initial setup of the OMR experiments.

## CRediT Statement

Thomas Rüland (Conceptualization, Data curation, Formal analysis, Investigation, Methodology, Software, Visualization, Writing - original draft)

Kerstin Doerenkamp (Investigation) Peter Severin Graff (Investigation) Sophie Wetz (Investigation) Anoushka Jain (Software)

Gerion Nabbefeld (Software, Writing - review & editing) Jana Gehlen (Formal analysis, Investigation)

Sara RJ Gilissen (Formal analysis, Investigation, Visualization, Writing - review & editing) Lutgarde Arckens (Conceptualization, Resources, Writing - review & editing)

Simon Musall (Conceptualization, Investigation, Methodology, Resources, Software)

Frank Müller (Conceptualization, Funding acquisition, Project administration, Resources, Supervision, Writing - review & editing)

Björn M. Kampa (Conceptualization, Funding acquisition, Project administration, Resources, Supervision, Writing - review & editing)

