## Supplementary Figures for "Ongoing activation of visual cortex and superior colliculus in the *rd10* mouse model of retinitis pigmentosa"

### Supplementary Material

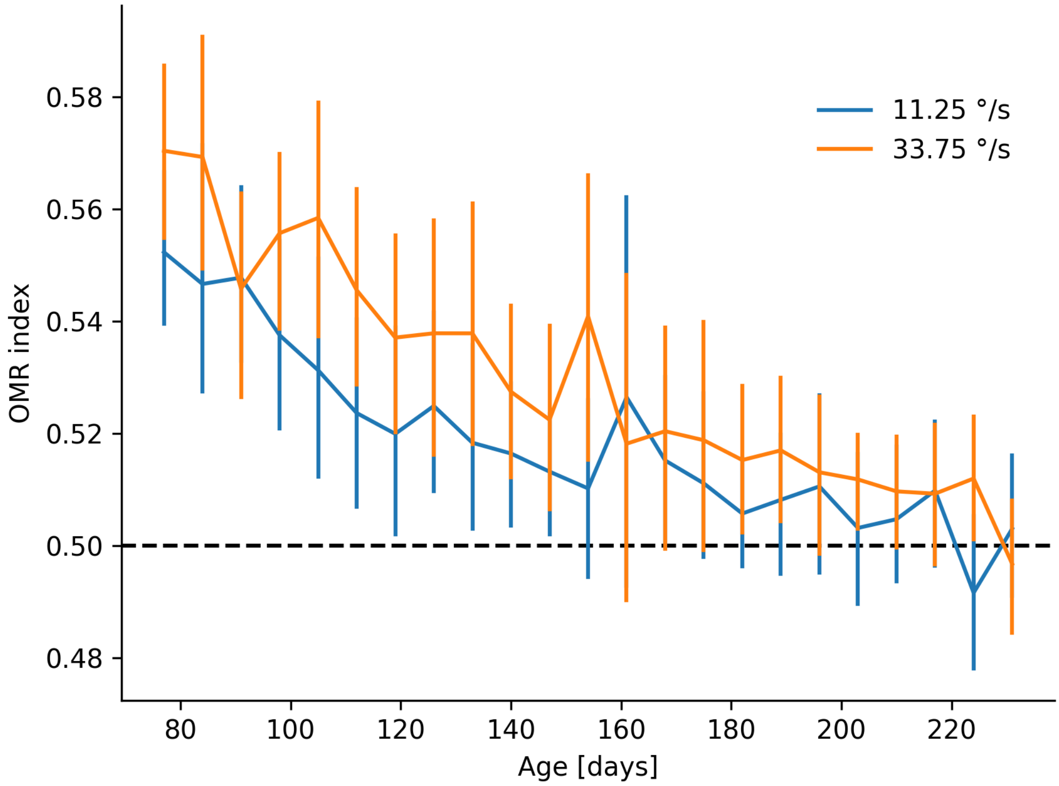


**Figure S1 Effect of grating speed on OMR index in *rd10* mice.** OMR index indicates grating following performance over time for grating speeds 11.25 °/s and 33.75 °/s in *rd10* mice at highest illumination strength (185 cd/m²). Errorbars depict the 95% confidence interval. Following performance does not significantly differ between both grating speeds across all time points measured. Differences measured using the Mann-Whitney U test, Bonferonni corrected ($p_{adjusted} \leq 0.05$) (*rd10*: $n_{mice}=3$, 11.25 °/s: $n_{sessions}=506$, 33.75 °/s: $n_{sessions}=520$).

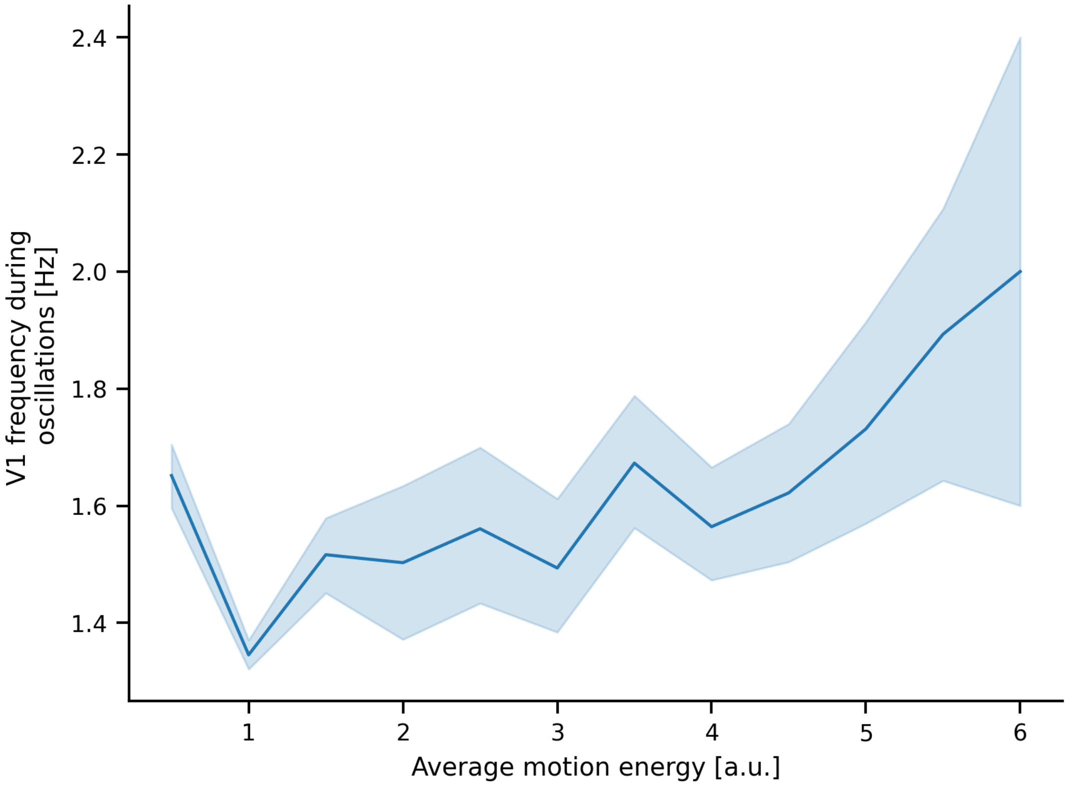


**Figure S2 Modulation of oscillation frequency in V1 by motion energy.** Shown is the average frequency with highest power spectral density in oscillation range (0.5 - 6Hz) during oscillation bouts of *rd10* recordings versus the average motion energy extracted from mice behavior videos. Shaded area depicts the 95% confidence interval. An increase in oscillation frequency with increasing motion energy can be observed. (*rd10*: $n_{mice}$=4, $n_{sessions}$=4).

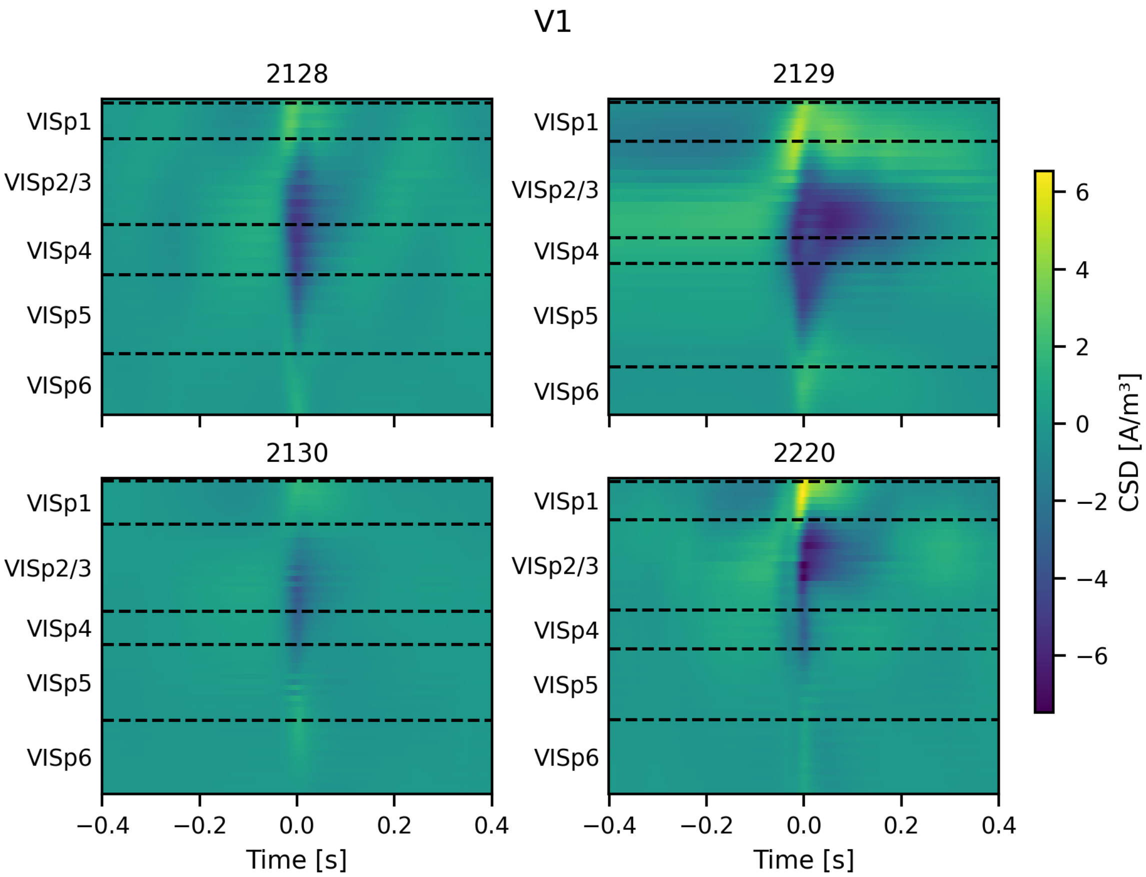


**Figure S3 Average current source density around oscillation events in V1.** Shown is the average current source density versus the depth in V1 400 ms before and after oscillation event occurrence for each oscillating session. Layers are indicated by dashed lines (*rd10*: $n_{mice}$=4, $n_{sessions}$=4).

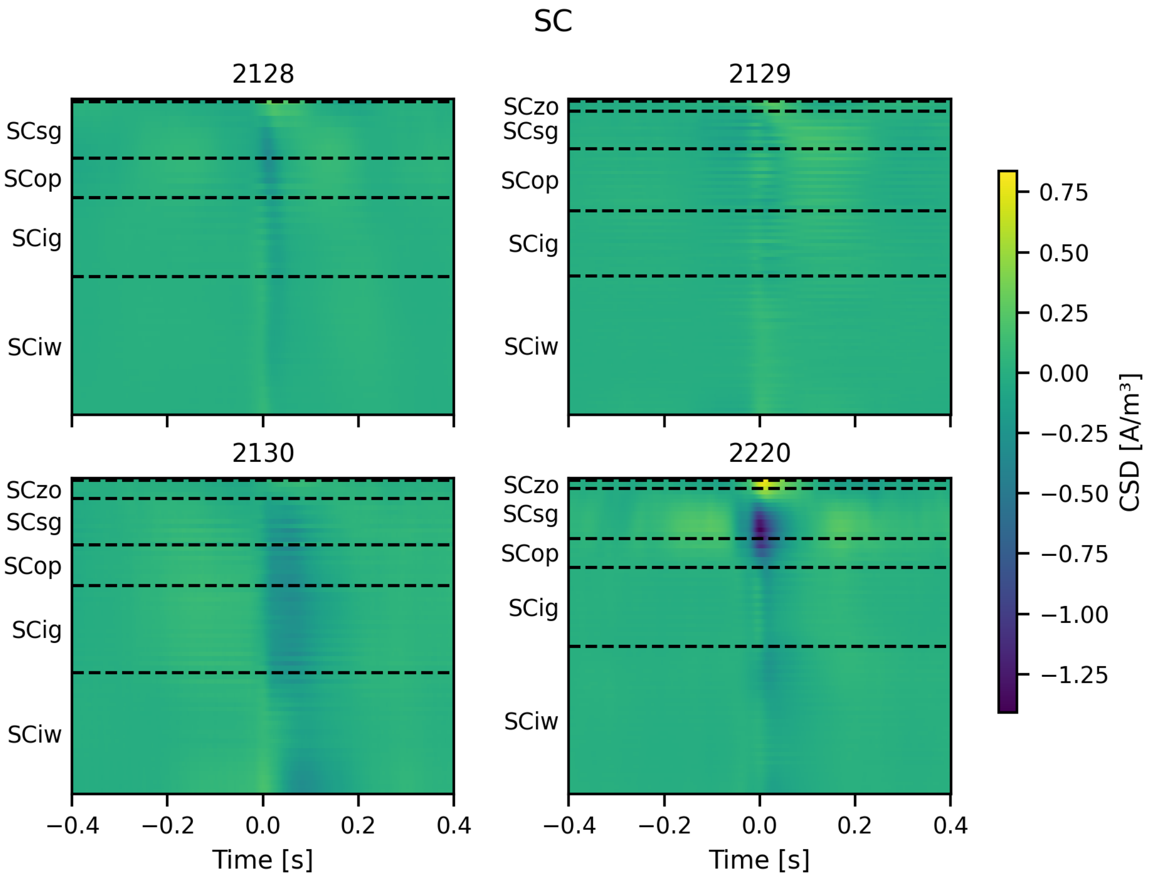


**Figure S4 Average current source density around oscillation events in SC.** Shown is the average current source density versus the depth in SC 400 ms before and after oscillation event occurrence for each oscillating session. Layers are indicated by dashed lines (*rd10*: $n_{mice}$=4, $n_{sessions}$=4).

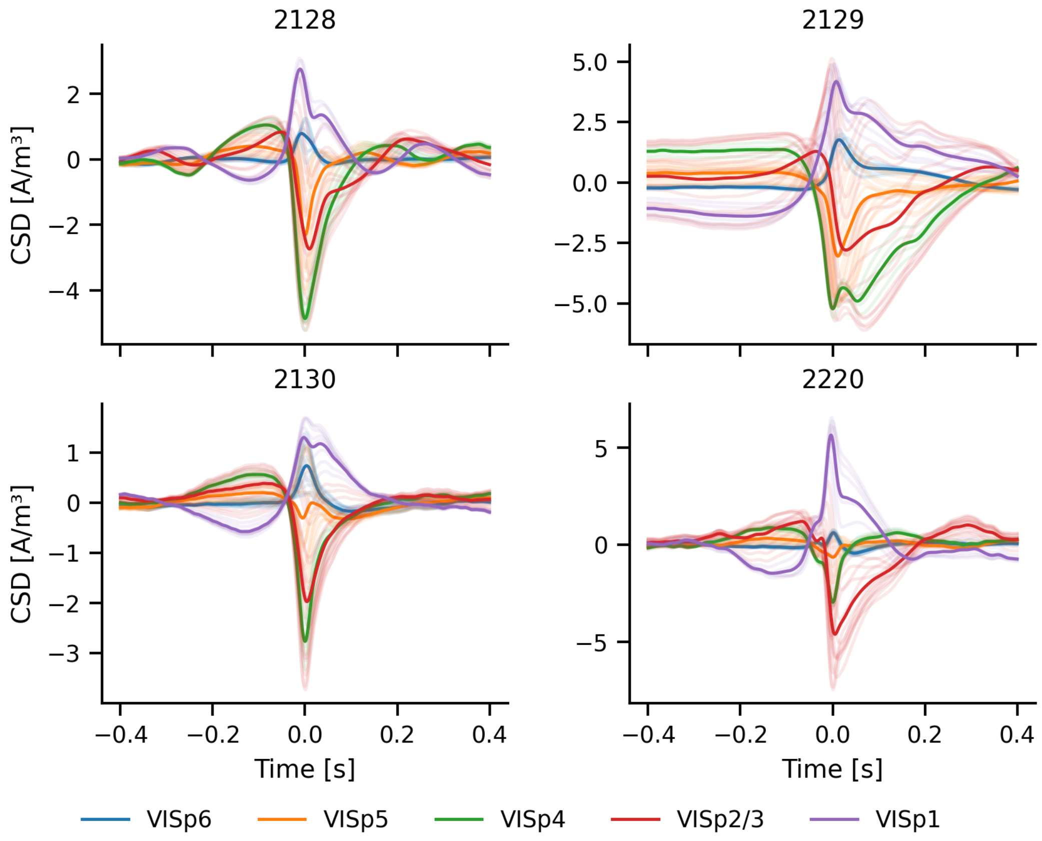


**Figure S5 Average current source density around oscillation events per layer in V1.** Shown is the average current source density per layer 400 ms before and after oscillation event occurrence for each oscillating session (*rd10*: $n_{mice}$=4, $n_{sessions}$=4). Faint lines show the individual channels for each area.

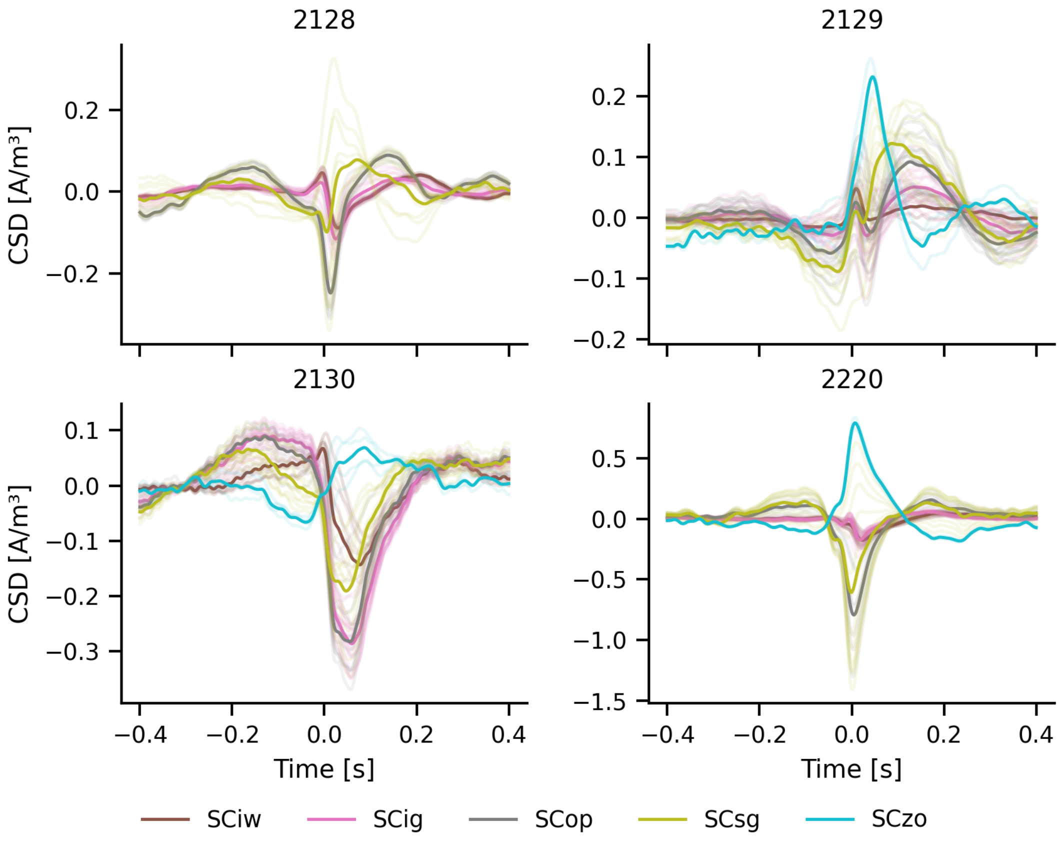


**Figure S6 Average current source density around oscillation events per layer in SC.** Shown is the average current source density per layer 400 ms before and after oscillation event occurrence for each oscillating session (*rd10*: $n_{mice}$=4, $n_{sessions}$=4). Faint lines show the individual channels for each area.

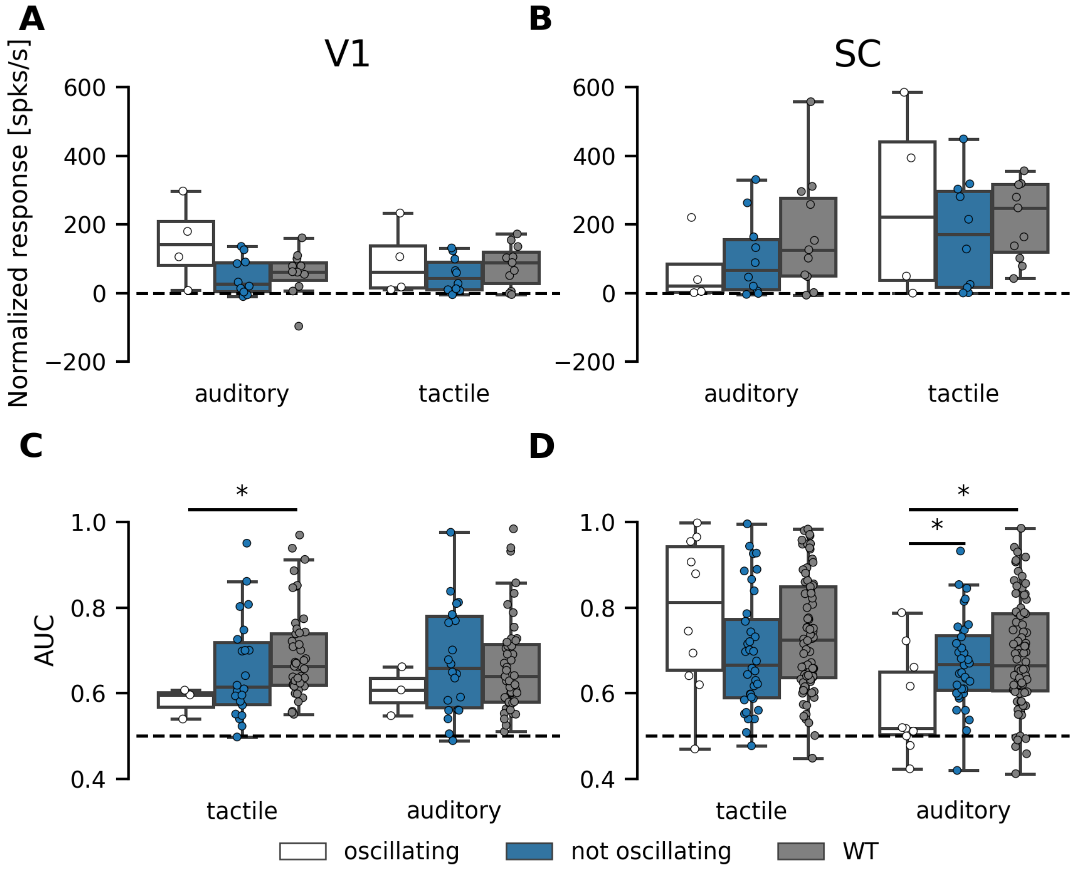


**Figure S7 Responses to non-visual stimulation in oscillating, non-oscillating and WT sessions. (A, B)** Average population responses for V1 (A) and SC (B) for oscillating (white) , non-oscillating (blue) and WT (gray) sessions. Differences were measured using the Mann-Whitney U test ($p \leq 0.05$) (*rd10* oscillating: $n_{mice}$=4, $n_{sessions}$=4, *rd10* not oscillating: $n_{mice}$=4, $n_{sessions}$=10, WT: $n_{mice}$=3, $n_{sessions}$=11,). **(C, D)** Non-visually responsive single-units in V1 (C) and SC (D) for oscillating (white) , non-oscillating (blue) and WT (gray) sessions. Differences measured using the Mann-Whitney U test ($p \leq 0.05$) (*rd10* oscillating: $n_{mice}$=4, $n_{sessions}$=4, V1: $n_{units}$=3, SC: $n_{units}$=10, *rd10* not oscillating: $n_{mice}$=4, $n_{sessions}$=10, V1: $n_{units}$=22, SC: $n_{units}$=36, WT: $n_{mice}$=3, $n_{sessions}$=11, V1: $n_{units}$=47, SC: $n_{units}$=80).

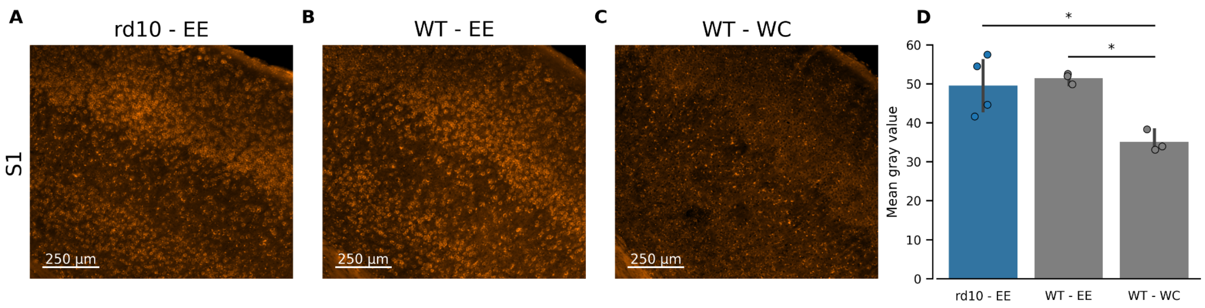


**Figure S8 Immediate early gene expression in response to tactile stimulation. (A-C)** Example field of views showing the somatosensory cortex of rd10 and WT mice after being exposed to an enriched environment without intact whiskers (A, B) and with cut whiskers (C). **(D)** Average gray value across all mice of all conditions shown in A-C. While no difference between WT and rd10 is visible in intact whisker mice, mice with cut whiskers show a decreased activity in the barrel field of somatosensory cortex despite exposure to an enriched environment. Differences were tested for differences using a One-Way Anova ($p \leq 0.05$) and a subsequent Tukey's multiple comparison test ($p_{adjusted} \leq 0.05$) (*rd10 - EE S1*: $n_{mice}=4$, S1 - EE: $n_{mice}=3$, S1 - WC: $n_{mice}=3$).
